# JuLI: accurate detection of DNA fusions in clinical sequencing for precision oncology

**DOI:** 10.1101/521039

**Authors:** Hyun-Tae Shin, Nayoung K. D. Kim, Jae Won Yun, Boram Lee, Sungkyu Kyung, Ki-Wook Lee, Daeun Ryu, Jinho Kim, Joon Seol Bae, Donghyun Park, Yoon-La Choi, Se-Hoon Lee, Myung-Ju Ahn, Keunchil Park, Woong-Yang Park

## Abstract

Accurate detection of genomic fusions by high-throughput sequencing in clinical samples with inadequate tumor purity and formalin-fixed paraffin embedded (FFPE) tissue is an essential task in precise oncology. We developed the fusion detection algorithm Junction Location Identifier (JuLI) for optimization of high-depth clinical sequencing. We implemented novel filtering steps to minimize false positives and a joint calling function to increase sensitivity in clinical setting. We comprehensively validated the algorithm using high-depth sequencing data from cancer cell lines and clinical samples and whole genome sequencing data from NA12878. We showed that JuLI outperformed state-of-the-art fusion callers in cases with high-depth clinical sequencing and rescued a driver fusion from false negative in plasma cell-free DNA. JuLI is freely available via GitHub (https://github.com/sgilab/JuLI).

## INTRODUCTION

High-throughput sequencing is becoming increasingly prevalent in precision cancer medicine worldwide. In the Republic of Korea and United States of America, assays using high-throughput sequencing have received regulatory approval as companion diagnostic tests for personalized care (http://www.fda.gov/MedicalDevices/ProductsandMedicalProcedures/InVitroDiagnostics/ucm330711.htm, http://www.mohw.go.kr/react/jb/sjb0406vw.jsp?PAR_MENU_ID=03&MENU_ID=030406&CONT_SEQ=338288&page=1). Most assays use sequencing technology to identify clinically actionable single nucleotide variants (SNVs) and small insertions/deletions (indels) because they are relatively easy to detect and interpret. However, some cancers such as *ALK*-rearranged non– small cell lung cancers (NSCLCs) and *BCR/ABL*-rearranged chronic myeloid leukemias (CMLs) are driven by somatic genomic fusions that cannot be detected by these methods for SNVs/indels. Patients with these oncogenic fusions respond to tyrosine kinase inhibitors (TKIs), and such genomic changes are now key therapeutic targets (Druker et al. 2001; Awad and Shaw 2014).

A number of factors are prerequisite for accurate detection of genomic fusions in the clinical setting. First, obtaining a representative specimen that provides an adequate amount of tumor sample for genome profiling is an ongoing challenge. Our previous study has shown that numerous important variants are present at a low allelic fraction (Shin et al. 2017). Unlike tissues used for research, tissues from clinical procedures, such as biopsies, tend to have inadequate tumor purity. Recently, cell-free DNA (cfDNA) testing by ultra-deep sequencing has been introduced for genotyping primary cancers and monitoring of post-treatment recurrence in oncology, and this test aims to detect approximately 0.1% of allele fractions (Oellerich et al. 2017; Phallen et al. 2017; Christensen et al. 2018). Furthermore, considering the heterogeneity of individual tumors, complete profiling of a tumor may require multiple samplings from different regions, which is not clinically feasible. To capture these low fraction variants, sufficient sequencing coverage and specialized algorithms are imperative for a clinical assay. Second, it is important to obtain a sufficient quality of formaldehyde-fixed paraffin-embedded (FFPE) specimens for genome profiling. FFPE is preferred for most molecular analyses of clinical pathologies because of its advantages in collection and storage. However, formalin fixation results in DNA and RNA damage, which is affected by various preanalytical factors, such as duration of storage, formalin fixation, and ischemic time (Evers et al. 2011; Spencer et al. 2013; Araujo et al. 2015). These fragmented nucleic acids act as noise and may make it difficult to detect oncogenic fusions. The detection of genomic fusions in clinical samples tends to be challenging because of the above-mentioned problems.

As the importance of detecting genomic fusions in clinical decision-making continues to increase, a critical area for improvement is currently the accuracy of detecting actionable fusions for the realization of precision cancer medicine. In the present study, we focused on improving the reliability of detecting somatic actionable fusions in cancer using high-depth DNA sequencing. To address the above problems, we developed a fusion detection algorithm optimized for clinical purposes and validated this algorithm using cancer cell lines with known driver fusions and **459** NSCLC samples with known *ALK* fusion and/or *RET* fusion status and **46** prostate cancer samples with known *TMPRSS2* fusion status (**Supplementary Table 1**).

## MATERIALS AND METHODS

### Study design

The Institutional Review Board (IRB) of Samsung Medical Center (SMC) approved this study. NSCLC samples were obtained at SMC between March 2014 and February 2017 with informed consent from some patients, whereas consent was waived by the IRB for others. The inclusion criteria for samples in this study were as follows: (i) sample was profiled using CancerSCAN™ (Shin et al. 2017) or LiquidSCAN™ (Park et al. 2018), the custom sequencing platforms of SMC; (ii) clinical information of the patient was stored in the clinical data warehouse of SMC.

### Panel design for fusion detection

Samples were prepared and analyzed using CancerSCAN™ or LiquidSCAN™, targeted-sequencing platforms designed at SMC (**Supplementary Table 1**) (Shin et al. 2017; Park et al. 2018). To identify fusions using a targeted panel, we tiled across the “hotspot” introns that contain well-known breakpoints of a set of clinically relevant fusions. Introns of five genes from an 83-gene panel (CancerSCAN version 1 and LiquidSCAN version 1) and introns of 22 genes from a 381-gene panel (CancerSCAN version 2) were densely covered with capture probes. All panels targeted hotspot introns of *ALK*. The average DNA fragment size of the platform was approximately 180 bp and the read length was 100 bp, thus, indicating that most fragments were fully sequenced. The other specific details of the panels can be found in previously reported papers (Shin et al. 2017; Park et al. 2018).

### Cell line mix experiment

Four cell lines (H2228, BHP10-3, U118MG, and SK-NEP-1) known to harbor specific fusions were used (**Supplementary Table 2**). The cell lines were cultured in our laboratory. Before extraction of DNA, the cells were washed two times with PBS. When the samples were pooled, the value from the Qubit HS assay (Life Technologies) was used, and DNAs were mixed equally to a total amount of 500 ng.

### PCR validation of fusions

The reference sequence of a target gene and breakpoint region was retrieved from the UCSC genome browser (http://genome.ucsc.edu/cgi-bin/hgBlat). A target-specific primer was designed using Primer3 for PCR on the basis of the reference sequence and was confirmed using Primer-BLAST (National Institutes of Health; NIH; **Supplementary Table 3**). The translocation target gene was amplified by PCR using specific primers. The cycling conditions were as follows: 94°C for 5 min, followed by 44 cycles of denaturation (94°C for 30 s), annealing (60°C for 1 min), and extension (72°C for 1 min), with final extension at 72°C for 10 min. The reactions were performed using HelixAmp TM Ready-2X-Go Hot-Taq (Nanohelix, Korea). Sequences of the PCR products were determined by an automated method (ABI Prism 3730) using the Big Dye Terminator Kit (Applied Biosystems, Foster City, CA, USA). Translocation breakpoint region sequences were verified by means of BLAST (NIH) and DNAstar (Lasergene).

### Alignment and preprocessing

Paired-end reads were aligned using BWA-MEM at its default settings (Li and Durbin 2009) with the human reference genome (hg19). Aligned reads with mapping quality <20 were filtered out, and the remaining reads were sorted using SAMtools (Li et al. 2009). To prepare appropriate input BAM files for other callers, we employed MarkDuplicates of Picard (Broad Institute), which is commonly used for marking and removal of duplicate reads (McKenna et al. 2010).

### Workflow for fusion identification

#### Fusion detection algorithm

To identify genomic fusions for clinical applications, we developed an algorithm called Junction Location Identifier (JuLI) with the aim of reducing the number of false positives generated while maintaining sensitivity. Initially, basic statistics of the BAM files, such as read length and median insert size, are calculated and used for further steps. Candidate breaks are then defined using two or more clipped reads, including at least one soft-clipped read, against the genome reference. If a matched normal sample is available as a control, breaks with twice the cutoff value of the clipped reads are scanned in the normal sample, and candidate breaks that overlapped with the breaks in the normal sample are excluded. If a set of normal samples is available, a control panel can be generated using a function in JuLI, which incorporates the breakpoints in multiple samples. All the samples in the present study were processed without matched normal or control panel filtering. The algorithm then involves two separate parts, viz., discordant and split read analyses. The user can set all parameters of each step.

#### Discordant read analysis

As JuLI does not remove duplicate reads as a part of the algorithm, counting supporting reads is very important to reduce the number of false positive calls. JuLI first uses information, including the genomic positions of both paired reads, CIGAR (Concise Idiosyncratic Gapped Alignment Report) strings, and the QNAME of sequencing reads in the BAM file, to reduce redundant duplicated or noise signals. Candidate breaks with fewer than three unique discordant reads are filtered. Next, consensus contigs from the matched and clipped side of each candidate break are generated. The average number of pairwise differences, representing nucleotide diversity (*π*), between the reads and the consensus contig on both sides of the candidate break is calculated as follows:

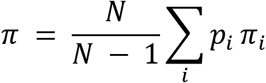

where *N* is the number of reads across the break, *ρ_i_* is the frequency of the *i*th read across the break, and *π_i_* is the proportion of bases that differ between the read and consensus contig truncated to the read length. If the normalized nucleotide diversity of either the clipped or matched side is higher than 2.0, the break is excluded from further processing. The normalized nucleotide diversity is calculated using the following formulae:

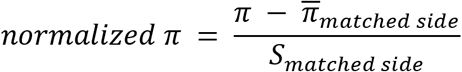

where 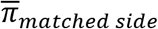 is the mean of nucleotide diversity of matched sides, and *S_matched side_* denotes the standard deviation of nucleotide diversity of the matched sides. Candidate breaks that pass the filters described above are paired with each other using the pair information, and split side contigs of each pair are aligned to the matched side contigs of their partners. If one of the two pairs matches more than 70% of the split contig length and is longer than 10 bp, the pair is called a fusion event. If there are no candidate pairs that passed the filters, a fusion event is defined if more than six discordant reads formed a cluster and more than 70% and more than 20 bp of the split contig is mapped to the reference sequence of the cluster region.

#### Split read analysis

Split read and discordant read analyses are conducted similarly. Candidate breaks with fewer than three split reads are filtered and then subjected to the following filtering steps, including nucleotide diversity analysis and pairwise local alignment. As JuLI is based on split information, fusions with a length less than half the read length are not considered. In split read analysis, if both pairs matched ≥70% of the length of the split contigs, the pair is considered a fusion event.

#### Joint call analysis

The joint call combines information from multiple BAM files in each analysis step and separates the numbers of each supporting read in the final step to produce individual results of the BAM files. If some fusion events have been previously defined in other BAM files, the fusions can be efficiently detected by specifying the target area using the BED (Browser Extensible Data) format. This is extremely useful for the case in cfDNA analysis as acquired serial samples for cfDNA may not have enough supporting reads, which makes it difficult to detect the events (see Discussion).

### Settings of algorithms

#### We carefully studied the documentation for each algorithm to determine and apply parameters that could be optimized in the clinical sample data

##### JuLI

All analyses were performed using JuLI v.0.1.3 with the default parameters. Fusion events in the UCSC gap database were excluded from further analysis.

##### SvABA

All analyses were performed using SvABA v 134 (Wala et al. 2018). We applied a -M flag so that the number of “weird reads” was not limited in highly fragmented FFPEs. We employed sorted, indexed, and duplication-free BAMs for SvABA. The command line for the analysis was as follows:

~~~
*$svaba run −t $INPUT.bam −p 1 -G $reference.fa −a sample_id −M 100000*
~~~

##### Delly

We used Delly v.0.7.8 (Rausch et al. 2012) for all analyses with the default parameters. We preprocessed BAM files as recommended by the developers (sorting, indexing, and duplicate marking). We applied the exclusion regions of the hg19 reference included in the Delly source code. The Delly command line for the analysis was as follows:

~~~
*$delly call -x human.hg19.excl.tsv -o $OUTPUT.bcf -g $reference.fa $INPUT.bam*
~~~

We converted the output with BCF (binary variant call format) to VCF (variant call format) using BCFtools, which was included as a submodule in Delly. We selected the results of VCF that passed the quality filter for all analyses.

##### Manta

All analyses were performed using Manta v.1.2.2 (Chen et al. 2016). We disabled all high-depth filters by applying the --exome flag during configuration for high-depth sequencing data. We analyzed sorted, indexed, and duplication-free BAMs using Manta. The command line for configuring was as follows:

~~~
*$configManta.py --tumorBam $INPUT.bam --referenceFasta $reference.fa --runDir
$OUTPUT_DIR --exome*
~~~

Next, we launched a workflow run script with a single node using the following command line for execution:

~~~
*$ OUTPUT_DIR/runWorkflow.py -m local -j 1*
~~~

We selected the results of VCF that passed the quality filter for all analyses. We applied the high-depth filter parameter for whole-genome sequencing (WGS) analysis.

##### LUMPY

We used LUMPY v.0.2.8 for all analyses (Layer et al. 2014). We analyzed sorted, indexed, and duplication-free BAM files for LUMPY. We split the BAM file into paired-end and split-read files using SAMtools v.0.1.19 (Li et al. 2009) with the recommended parameters and performed statistical analysis of the library sizes by means of a script in LUMPY. Then, the following LUMPY command line for fusion detection was executed:

~~~
*$-mw 4 -tt 0.0 – pe bam_file:$INPUT.discordant.pe.bam,histo_file:$INPUT.pe.histo,mean:$MEAN,stdev:$STDEV,r
ead_length:100,min_non_overlap:100,discordant_z:4,back_distance:20,weight:1,id:1,min_map
ping_threshold:20 -sr
bam_file:$INPUT.sr.bam,back_distance:20,weight:1,id:2,min_mapping_threshold:20 >
$OUTPUT.pesr.bedpe*
~~~

##### novoBreak

All analyses were performed using novoBreak v 1.1(Chong et al. 2017). We employed sorted, indexed, and duplication-free BAMs for novoBreak. We simulated a control BAM file using wgsim (H. 2011) and used the output as control input to novoBreak. The command line for the analysis was as follows:

~~~
*$run_novoBreak.sh $novoBreak_exe_dir $reference.fa $INPUT.bam $CONTROL.bam 1
$OUTPUT_DIR*
~~~

## RESULTS

### Development of a fusion detection algorithm for clinical sequencing

Since 2014, we have used a custom-designed panel (CancerSCAN™) for precision oncology that covers up to 381 cancer-related genes, including introns containing frequent breakpoints in selected fusion genes (Shin et al. 2017). To obtain high detection rates, we ensured a mean sequencing coverage of approximately 1200X and a target insert size of approximately 180bp in the initial alignment. We have developed several algorithms to improve the accuracy of our platform. For fusion detection, here, we developed JuLI, which was optimized for high-depth sequencing (**Fig. 1a–d**). JuLI utilizes information from both discordant and proper pair reads to detect a wide range of structural variations (SVs), including duplications, deletions, inversions, and interchromosomal translocations, at single-nucleotide resolution. Generally, it is preferable to conduct high-depth sequencing with relatively short insert sizes (150–200 bp) to achieve high sensitivity of target-enriched sequencing in various platforms, including panel-based platforms. However, PCR duplicates generated during preprocessing for sequencing may result in overestimation of variants, and this situation may cause false positive results that could be even worse with short insert sizes (Zhou et al. 2014). To avoid this problem, identifying duplicates using Picard (McKenna et al. 2010) or SAMtools (Li et al. 2009) is a necessary step in general bioinformatics analysis. However, because this process uses only limited information on sequence alignment map (SAM) files, it is possible to unintentionally remove reads with evidence of rearrangement (**fig. S1**), which may, thus, affect the sensitivity of detecting lower tumor cell content. We carefully counted reads supporting candidate breaks by determining duplicate fragments using CIGAR and pair locations without applying a general deduplication step (**Fig. 1b**; see **Methods**). Next, the candidates with sufficient supporting reads were subjected to the following two filtering steps. First, we measured nucleotide diversity (*π*), which is the average number of pairwise differences between the reads and the consensus contig, and the breaks with high nucleotide diversity were excluded from further processing (**Fig. 1c**; see **Methods**). Second, the candidate break and partner breaks were paired via pair information and compared by pairwise local alignments **(Fig. 1d**; see **Methods)**. Through these filtering steps, we were able to accurately detect fusions by reducing the number of false positives.

**Fig. 1.**
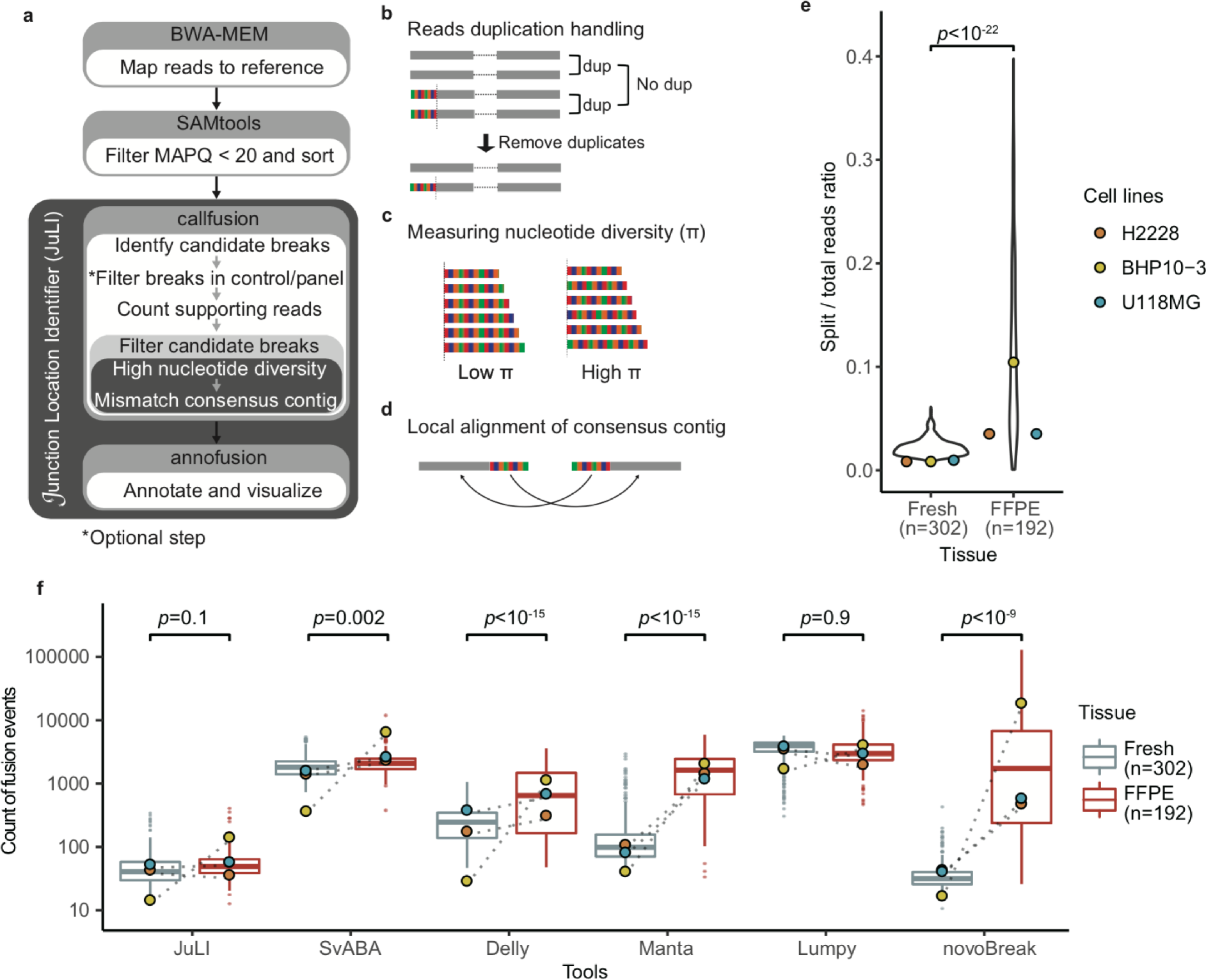
The fusion detection algorithm for clinical sequencing. **(a)** The scheme of Junction Location Identifier (JuLI). JuLI implements novel filtering steps to reduce the number of false positives while maintaining sensitivity by fine-tuning the counting of supporting reads without duplicate removal. **(b)** JuLI uses information, including the genomic positions, CIGAR strings, and read names in the BAM file, to reduce redundant duplicated or noise signals. **(c)** After measuring the nucleotide diversity of the breaks, JuLI filters breaks with high nucleotide diversity for the analysis. **(d)** The candidate breaks are paired with each other using pair information, and split side contigs of each pair are aligned to the matched side contigs of their partners. **(e)** The ratios of split read to the total read counts for formalin-fixed paraffin-embedded (FFPE), fresh clinical tissue samples (n = 494), and pair cell lines (n = 3) tested by CancerSCAN™. For FFPE samples, split reads and variability of split reads increased significantly (*t* test, *p* < 10^−22^). **(f)** Variant counts obtained from the callers in patient samples (n = 494) and pair cell lines (n = 3). Note that some callers showed increasing variant counts in FFPE tissues. This phenomenon was due to the low quality of FFPE samples because of DNA degradation or damage.

### Effects of damaged DNA in FFPE tissues

As mentioned above, one of the challenges in analyzing clinical samples is that FFPE tissues usually contain degraded DNA and smaller fragment sizes (Spencer et al. 2013). As a consequence, the ratio of split to total reads is substantially higher in FFPE samples than that in fresh samples examined by CancerSCAN (*t* test, *p* < 10^−22^; **Fig. 1e**). To eliminate the differences between individual samples, three pairs of fresh and routinely processed FFPE cancer cell lines were chosen for sequencing to compare tissue effects. Furthermore, differences in the split to total read ratio were also observed (**Fig. 1e**). An increase in the numbers of split reads could affect noise in fusion analyses and may cause numerous false positive events. To compare FFPE effects and for further analysis, we chose several state-of-the-art fusion callers, including SvABA (Wala et al. 2018), Delly (Rausch et al. 2012), Manta (Chen et al. 2016), LUMPY (Layer et al. 2014), and novoBreak (Chong et al. 2017), that use split and discordant read information, similar to JuLI. In the comparison of fusion events count, we observed a significant increase in count of fusion events in FFPE tissues when using SvABA, Delly, Manta, LUMPY, and novoBreak (**Fig. 1f**). The count of fusion events of JuLI and LUMPY was not affected by the tissue type, but the count of LUMPY was ten-times higher than that of JuLI, regardless of the tissue type (**Fig. 1f**). Analysis of three paired fresh and routine FFPE cancer cell line specimens revealed differences in counts of fusion events between the FFPE and fresh specimens. BHP10-3 revealed the highest change in the split/total read ratio (**Fig. 1e**) and showed the highest difference in fusion counts using most callers (**Fig. 1f**). Numerous split reads were observed in the FFPE specimen of BHP10-3 probably because of DNA damage during sample preparation (**fig. S2**). For the tools affected by FFPE tissues, the count of fusion events was positively correlated with split/total reads ratio (**fig. S3)**. However, JuLI showed the least increase in the number of fusion events with increasing split/total reads ratio (**fig. S3)**. Low quality of FFPE tissue can cause numerous false positive results with most callers, but such quality issues did not significantly affect the results yielded by JuLI.

### Validation of analytical sensitivity on cancer cell lines and patient’s samples

To evaluate the accuracy of the algorithm over a wide range of tumor purity, we adopted experimental schemes designed by Frampton et al. (Frampton et al. 2013). To simulate different tumor purity levels, four cancer cell lines harboring known fusions and a normal sample were manually mixed at different ratios, generating a range of expected tumor purity levels (5%–100%) (**Supplementary Table 2**). All cell line specimens were profiled using CancerSCAN™ version 1, which targeted 83 genes. The mixed fraction of the fusions showed a high correlation (correlation coefficient [*r*] = 0.95) with the relative value of the normalized supporting reads (**fig. S4**). We observed that JuLI, SvABA, Delly, Manta, and LUMPY achieved 100% sensitivity (32/32), but novoBreak missed one large deletion between *GOPC* and *ROS1* with 5% mix fraction in this experiment (**Supplementary Table 4**). In addition, 37 fusions in patients’ tissues detected by JuLI with a wide range of supporting reads (range, 6–283) were validated by PCR to verify the estimated fusion breakpoints. The locations of all fusion sequences at the estimated breakpoints were confirmed (**Supplementary Table 3**).

### Performance validation using clinical samples

Because of the differences in the performance between callers depending on the range of fusion length (Wala et al. 2018), we measured the F1 score [the harmonic average of positive predictive value (PPV; also known as precision) and sensitivity (also known as recall)] of the callers according to the minimum fusion length in 494 clinical samples examined by CancerSCAN (**Fig. 2a and Supplementary Table 1**). Fusion results that are shorter than the minimum length in each caller were excluded from the comparison and the performance comparison criteria are described in the following paragraph. In all ranges of minimum fusion size, we observed that JuLI outperformed other callers. Although JuLI and SvABA were less affected by performance over the range of fusion sizes, Delly exhibited increased performance at a relatively long length of fusion. We observed that Manta, LUMPY, and novoBreak tended to have lower PPV compared to sensitivity (**Supplementary Table 5**) and a decrease in performance in predominantly FFPE tissues compared to that in fresh tissues (**Supplementary Table 5**). The minimum length of F1 score saturation for each caller was 800 bp for JuLI and SvABA, 1500 bp for Delly, 1900 bp for Manta, 1200 bp for LUMPY, and 1300 bp for novoBreak. In order to compare except for the regions with different performance, we compared the results except for the fusions with the length shorter than 1250bp, which is the median value of the performance saturation length of each caller.

**Fig. 2.**
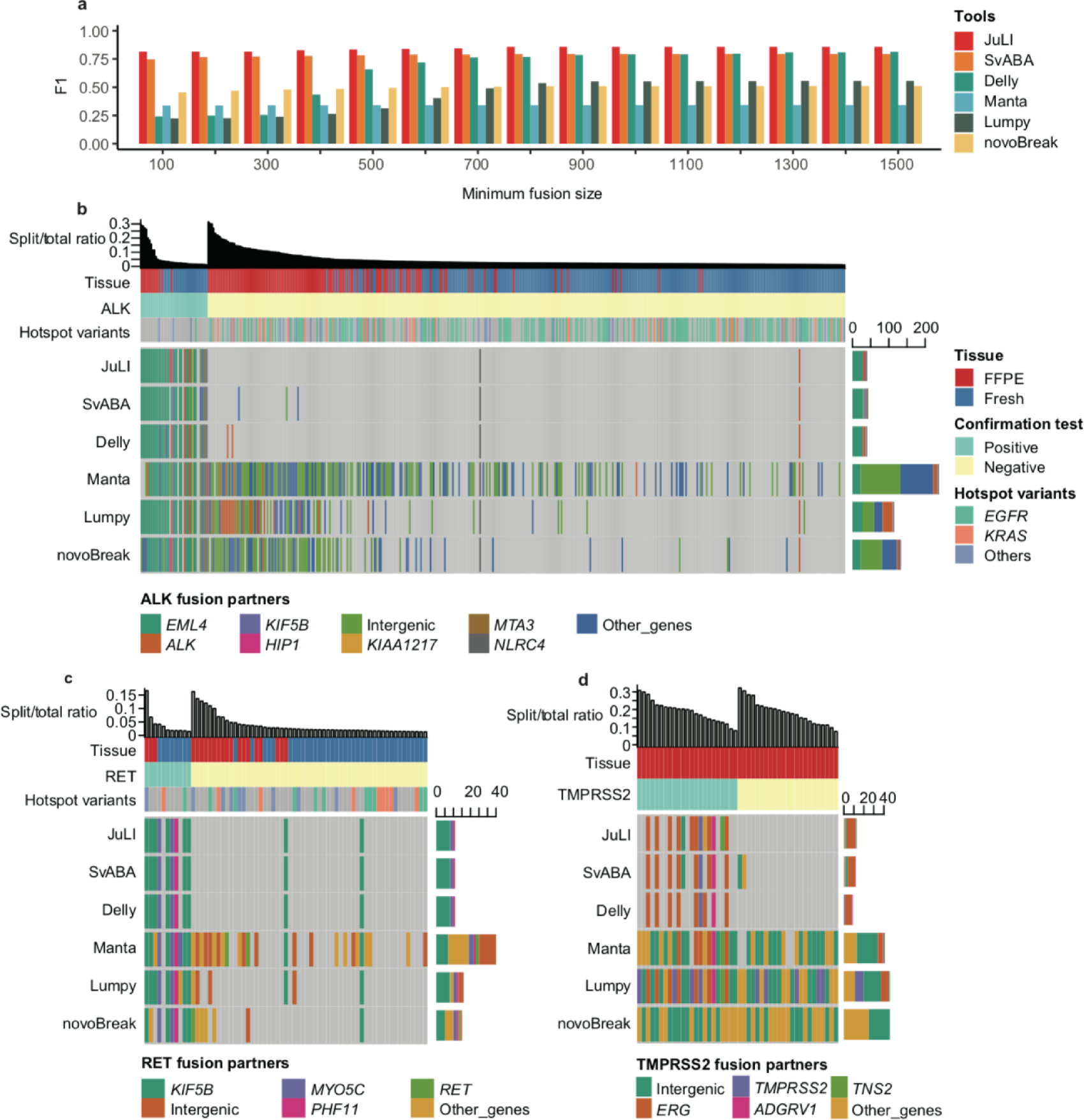
Validation on high-depth clinical samples. **(a)** The F1 score [the harmonic average of positive predictive value (PPV; also known as precision) and sensitivity (also known as recall)] of the callers according to the minimum fusion length in 494 clinical samples examined by CancerSCAN™. **(b)** Validation on 441 non-small cell lung cancer (NSCLC) samples with known *ALK* fusion status via IHC and/or FISH analyses. **(c)** Validation on 67 NSCLC samples with known *RET* fusion status via IHC and/or FISH analyses. **(d)** Validation on 46 prostate cancer samples with known *ERG* fusion status via IHC and/or FISH analyses. All samples were sequenced using Illumina Hi-Seq with high coverage (~ 1300X). When two or more events in the fusion gene were detected, we defined fusion on the basis of the most supportive read counts. Hotspot mutations of NSCLCs were in *EGFR* (L858R or exon 19 indel) or *KRAS* (G12, G13, or Q61). IHC, immunohistochemistry; FISH, fluorescence *in situ* hybridization; FFPE, formalin-fixed paraffin-embedded.

Activation of kinase gene by chromosomal rearrangement has been identified as a recurrent driver event in NSCLCs (Takeuchi et al. 2012; Pan et al. 2014). *ALK* rearrangement acts as an oncogenic driver in 4%–6% of NSCLCs (Takeuchi et al. 2012). In *ALK*-rearranged NSCLCs, ALK inhibitor demonstrates therapeutic efficacy in terms of improved survival, and the *EML4/ALK* variants and *ALK*-fusion partners may affect sensitivity to ALK inhibitors (Kwak et al. 2010; Shaw et al. 2013; Noh et al. 2017). *RET* rearrangements have been identified in 1%–2% of NSCLCs and are the potential therapeutic targets of multi-targeted kinase inhibitors (Pan et al. 2014; Lee et al. 2015). Therefore, accurate detection of an oncogenic fusion is important for clinical decision-making. Over the last four years, CancerSCAN™ has been used at the oncology clinic of SMC. We conducted performance validation in a prospective cohort of **448** patients with NSCLC and profiled *ALK* and/or *RET* status by immunohistochemistry (IHC) and/or fluorescence *in situ* hybridization (FISH). Of the 441 patients tested for ALK, 9.5% (42/441) were positive, and 67 patients were tested for RET, of which 16.4% (11/67) were positive (**Supplementary Table 1**). No patient was both *ALK*-and *RET*-positive, and the results of the IHC/FISH of *ALK* and other hotspot mutations in *EGFR* (L858R or exon 19indel) or *KRAS* (G12, G13, or Q61) showed a mutually exclusive pattern (Fisher’s exact test, *p* < 10−11). A total of **79** patients were profiled using CancerSCAN™ version 1, which targeted 83 genes, whereas the rest were profiled using version 2, which targeted 381 genes (Shin et al. 2017). Both V1 and V2 panels covered the same hotspot introns involved in *ALK* and *RET* rearrangement (introns of *ALK* between exons 19–21 and *RET* between exons 6–12).

As mentioned above, we considered fusion events that were ≥1250 bp in size, and ≥1 breaks were found in the analysis of *ALK* and *RET* region. Most *ALK* and *RET* activation cases involved the rearrangement or activating mutations that activate the kinase domain; in case of NSCLC, *ALK* and *RET* are primarily activated by fusion with various partners (Hallberg and Palmer 2013; Lee et al. 2015; Noh et al. 2017). Therefore, we assumed intragenic rearrangements in *ALK* and *RET* as a false positive. The respective sensitivity and PPV of *ALK* fusions were as follows: JuLI, 90.4% (38/42 samples) and 95.0% (38/40); SvABA, 88.0% (37/42) and 88.0% (37/42); Delly, 88.0% (37/42) and 90.2% (37/41); Manta, 83.3% (35/42) and 14.7% (35/238); LUMPY, 88.0% (37/42) and 32.3% (37/115); and novoBreak, 90.4% (38/42) and 28.6% (38/133) (**Fig. 2b)**. For *RET* fusions, JuLI, SvABA, and Delly achieved same sensitivity and PPV [81.8% (9/11 samples) and 81.8% (9/11), respectively]. The sensitivity and PPV of remaining callers were as follows: Manta, 90.9% (10/11) and 28.6% (10/35); LUMPY, 90.9% (10/11) and 62.5% (10/16); and novoBreak, 72.7% (8/11) and 53.3% (8/15) (**Fig. 2c)**. Six samples that yielded false negative results of *ALK* and *RET* in JuLI analysis also tested negative in most callers, and the tumor purity of these samples was significantly lower than that of the test-positive samples (**fig. S5**). Therefore, some false negatives may be due to low tumor purity. Four false positives of *ALK* and *RET* identified in JuLI results were observed in all other callers, and the fusions were clearly identified in browser view (**fig. S6**).

To further compare other clinically significant fusions, we retrospectively collected 46 archived prostate cancer samples and performed analysis of *ERG* fusion status by IHC and/or FISH. Twenty-three of the 46 patients (50.0%) were *ERG* fusion-positive (**Supplementary Table 1**). All patients with prostate cancer were profiled using CancerSCAN™ version 1, and the panel covered the hotspot introns between exons 1–6 of *TMPRSS2*, the most common fusion partner of *ERG* fusion (Barros-Silva et al. 2013). We measured the performance of the callers with the same criteria as those of NSCLC. The respective sensitivity and PPV of *ERG* fusions were as follows: JuLI, 56.5% (13/23 samples) and 100.0% (13/13); SvABA, 43.5% (10/23) and 83.3% (10/12); Delly, 39.1% (9/23) and 100.0% (9/9); Manta, 95.7% (22/23) and 53.7% (22/41); LUMPY, 100.0% (23/23) and 50.0% (23/46); and novoBreak, 100.0% (23/23) and 50.0% (23/46) **(Fig. 2d)**. There was no difference in purity distribution between true positive and false negative of JuLI. The relatively low sensitivity of this retrospective set may be due to other partners of *ERG* that were not targeted (Cancer Genome Atlas Research 2015). Overall, the number of false calls occurred as the split/total read ratio increased, but this issue had less effect in JuLI.

### Performance based on sequencing coverage

For panel-based high-throughput sequencing in clinical practice, test performance must at least be comparable to conventional molecular tests. The factors that constitute sufficient sequencing depth are influenced by tumor purity and clonality of variants as well as other characteristics of a patient’s tumor sample, including tissue preparation methods and sequencing platforms used. Lowering tumor purity reduces detection sensitivity by proportionally reducing the effective range of mutant alleles in tumor cells. In contrast to research samples, requirements for sufficient tumor purity for clinical specimens may not be met; therefore, it is important to have adequate coverage (Shin et al. 2017).

We conducted *in silico* down-sampling experiments (three iterations) using the *ALK* set of NSCLCs as an alternative method for investigating the effect of sequencing depth on performance. In down-sampling experiments, we observed that the average sensitivity of all the callers improved with increased coverage (**Fig. 3a** and **3c**). Although sensitivity increased at high coverage, PPV and specificity decreased in Manta, LUMPY, and novoBreak. In contrast, we noticed that JuLI, SvABA, and Delly were less affected by coverage. The F1 score improved in the 50–200× range, which is the typical range used in WGS or whole-exome sequencing (WES) in all callers (**Fig. 3b**). By contrast, the F1 score worsened at high depth (1340×) in Manta, LUMPY, and novoBreak, but not in others. Accuracy, which is the proportion of true results (both true positives and true negatives) among all the results, was also maintained at a high-depth range in JuLI, SvABA, and Delly in contrast to the other callers (**Fig. 3d**). Thus, we confirmed that sensitivity improved with increased sequencing depth; however, accuracy may decrease owing to increased noise levels above the threshold in some algorithms. To effectively apply high-throughput sequencing in a clinical setting, it is necessary t reduce such noise.

**Fig. 3.**
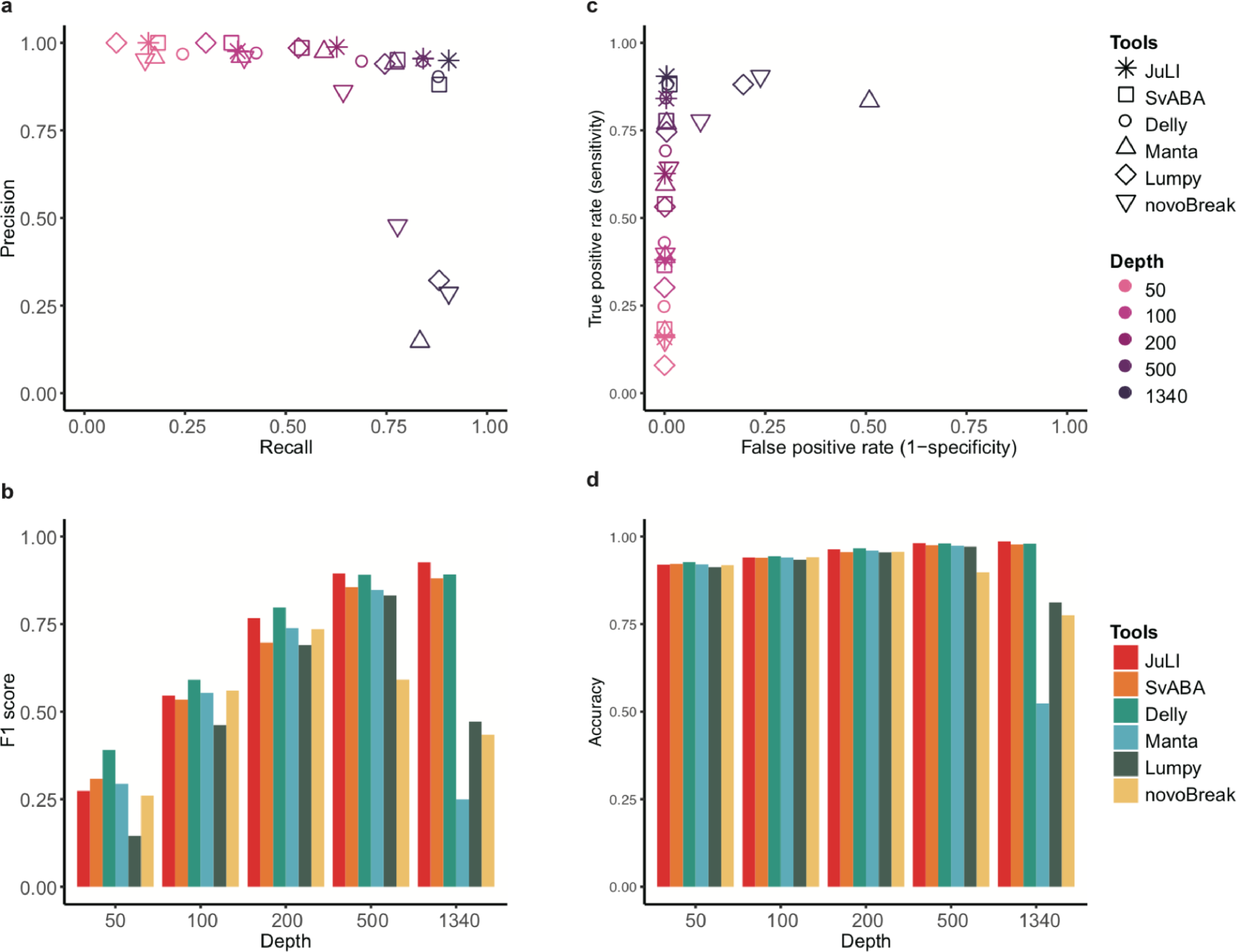
The effect of depth on fusion detection in clinical samples. **(a)** Positive predictive value (PPV, also known as precision) and sensitivity (also known as recall) based on *in silico* down-sampling experiments. A total of 441 non-small cell lung cancer (NSCLC) samples were down-sampled from the original depth (1340X), and the average performance was measured at each depth (three iterations). **(b)** The F1 score, which is the harmonic average of PPV and sensitivity, on the basis of the coverage change. **(c)** The receiver operating characteristic (ROC) curve. **(d)** Accuracy, which is the proportion of true results (both true positives and true negatives), among all the results. Error bars denote standard error of the mean.

### Sensitivity validation using WGS data

To estimate the sensitivity based on real WGS data, we downloaded raw FASTQ data of NA12878 from European Nucleotide Archive (ERA172924, https://www.ebi.ac.uk/ena/data/view/PRJEB3381). These data represent approximately 50X coverage, which has been widely used by tools for the estimation of a variety of variation tools. We compared results made by each tool to the truth set by Layer et al. (Layer et al. 2014), who developed LUMPY in 2014. They provided a truth set containing 4,095 deletions detected by at least one tool in the 50X dataset that were validated by split-read mapping analysis of independent long-read sequencing data from PacBio or Illumina platforms. In this comparison, LUMPY (47.4%; 1942/4095) was the most sensitive, followed by JuLI (41.9%; 1717/4095), SvABA (41.9%; 1716/4095), Delly (38.5%; 1575/4095), Manta (37.9%; 1552/4095), and novoBreak (37.7%; 1542/4095). We were able to confirm that the performance of JuLI was maintained as much as other callers even at a low depth, such as WGS; however, specificity or PPV representing the frequency of false positive calls was not evaluated due to the lack of true negative reference.

### Joint call to detect fusions with insufficient evidence

Recent reports have shown that the detection of *ALK* fusion in cfDNA is feasible in clinic setting (Paweletz et al. 2016; Thompson et al. 2016). Serial tumor sampling on progression has been helpful in determining the optimal subsequent treatment decision-making for patients. However, this is often complicated by insufficient tumor purity for molecular analysis and tumor heterogeneity (Dagogo-Jack et al. 2018). If a more sensitive detection is possible in a series of samples with insufficient supporting reads, slightly earlier decision-making can be made for precise medicine. To achieve more sensitive fusion detection in clinical sequencing, we implemented the joint call function in JuLI that can detect fusions with low supporting evidence in serial/multi-region sampling tissues (**Fig. 4a**). To verify the performance of this function, *in-silico* down-sampling experiments (mean coverage: 1X, 5X, and 10X with 100 iterations) were performed on mixed cell lines with relatively low cell ratios of 5-40% (**Supplementary Table 4**). In this simulation, most callers showed up to 1-2% sensitivity at 10X, joint call showed 40.6% sensitivity at 10X and could detect 7.3% even at 1X (**Supplementary Table 6**).

**Fig. 4.**
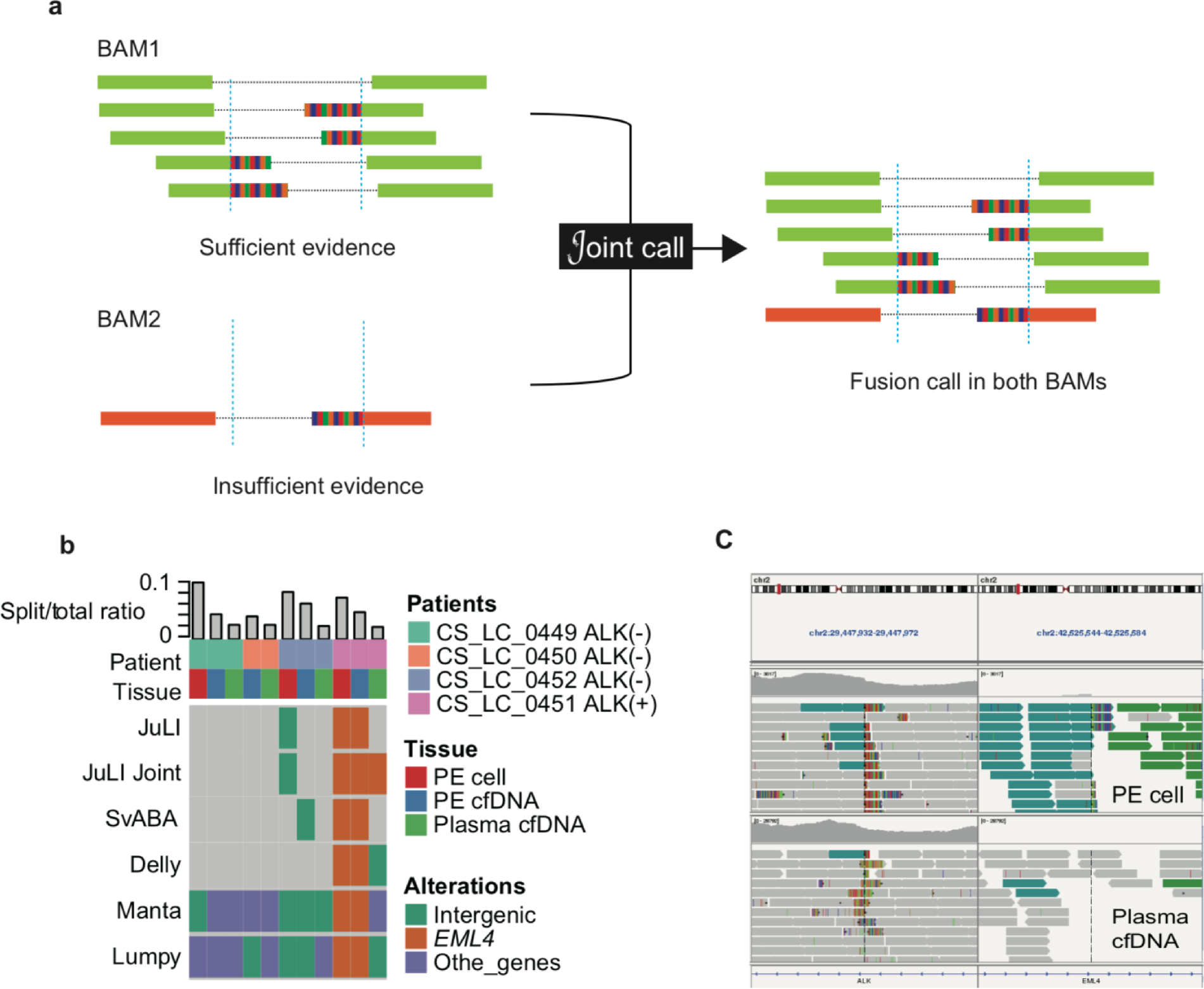
The joint call function to detect fusions with low supporting evidence in serial/multi-region sampling tissues. **(a)** The joint call function combines information from multiple BAM files and produces the individual result of the BAM files. **(b)** JuLI with the joint call function rescued the *EML4/ALK* fusion of CS_LC_0451 from false negative in plasma cell-free DNA. **(c)** Only two discordant reads supporting the *ALK* fusion were observed in plasma cfDNA of CS_LC_0451.

To confirm the utility of the joint call function of JuLI in clinic, we applied it to *ALK* detection in cfDNA of NSCLCs. Pleural effusion (PE) and peripheral plasma were collected from four patients with NSCLS, whose *ALK* status was confirmed in primary tissue (one positive and three negative; **Supplementary Table 1**). DNA of cells in PE, cfDNA of PE, and cfDNA of plasma were processed using LiquidSCAN™ (average coverage, approximately 4300X) and analyzed using the fusion callers. We excluded novoBreak from this analysis because some samples did not show any results in the ultra high-depth data. In this analysis, the callers, except JuLI, missed the *EML4/ALK* fusion in plasma cfDNA of the *ALK*-positive patient (CS_LC_0451), but JuLI was able to identify the fusion using the joint call function (**Fig. 4b**). There were only two discordant reads supporting the *ALK* fusion in plasma cfDNA of CS_LC_0451 (**Fig. 4c**).

### Annotation and visualization

Accurate annotation of rearrangements is critical for clinical decision-making. JuLI annotates functional consequences of genomic fusions that are identified using high-throughput sequencing data in a strand-specific manner (**fig. S7**). Even with breaks at the same location, this annotation approach allows the user to easily distinguish between positive and negative strand events. Moreover, JuLI provides three useful pieces of information. First, JuLI predicts whether the fusion transcript is in-frame or out-of-frame by means of the UCSC database (Kent et al. 2002). Second, JuLI provides the frequency of fusion events based on cancer types in COSMIC (Forbes et al. 2015). Third, JuLI annotates chimera protein domains via the UniProt (Apweiler et al. 2004) and Pfam databases (Finn et al. 2010). Graphically visualized fusion diagrams were automatically generated in PDF format showing all annotation results (**Fig. 5**).

**Fig. 5.**
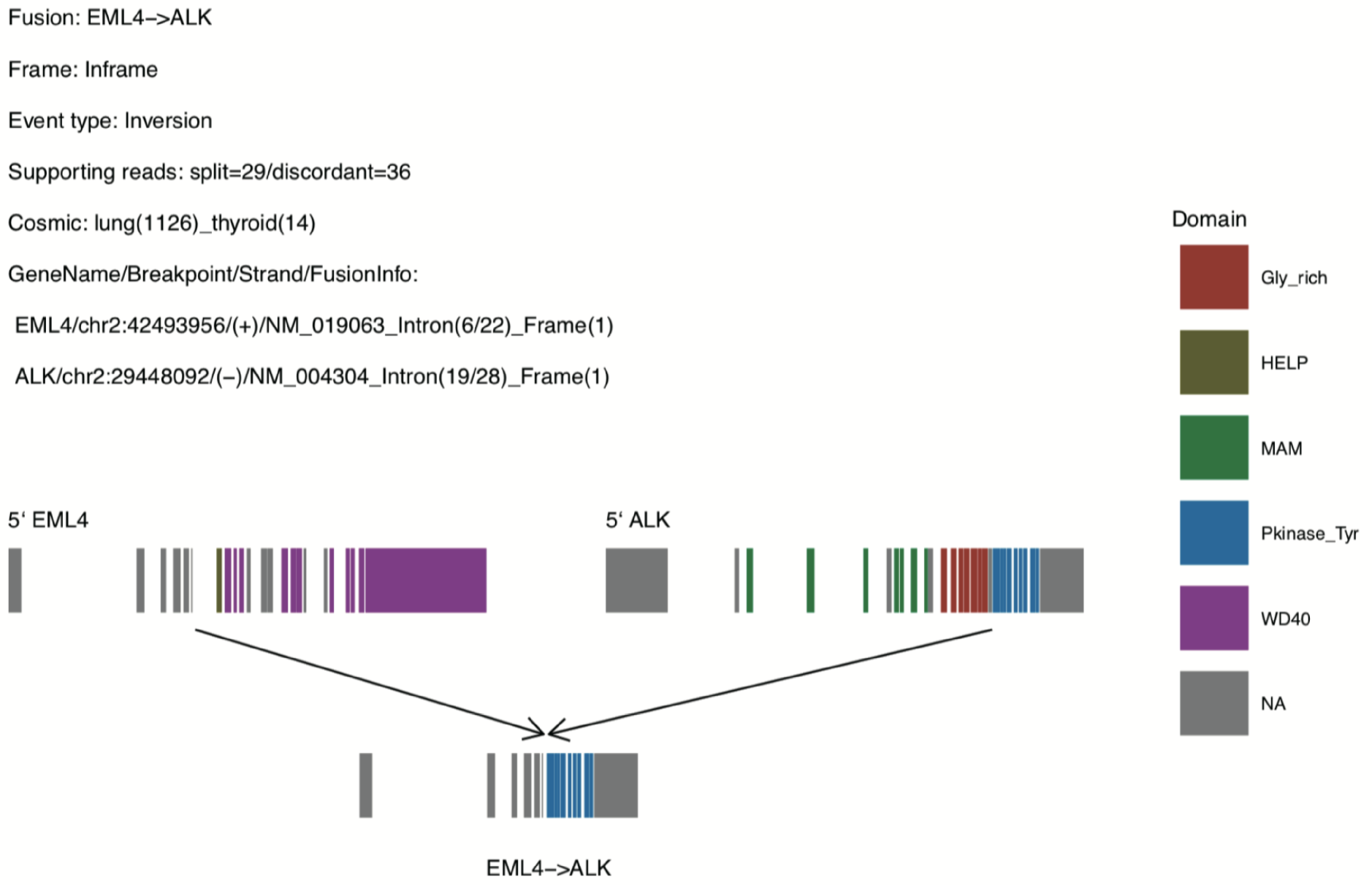
A representation showing Junction Location Identifier (JuLI) output with annotation and visualization. Annotation of fusions and a graphically visualized fusion diagram with the domain status in PDF format.

### Comparison of running time between paired FFPE and fresh tissues

We generated sequencing data from paired FFPE and fresh cancer cell lines using CancerSCAN V1, and of these cell lines, BHP10-3 showed the highest change in the split/total read ratio (**Fig. 1e**). To compare elapsed time, we measured BHP10-3 pair analysis time of each caller with 10 iterations (**fig. S8**). We observed a 1.1-to 29.3-fold increase in analysis time in low-quality FFPE tissue with these callers. Although JuLI was relatively slow in low quality FFPE tissue because it implements several steps to improve accuracy, the speed can be increased through parallel processing across multiple cores.

## DISCUSSION

To implement precision medicine at SMC, we developed JuLI, a novel fusion detection algorithm optimized for clinical application. We validated the tool on four cancer cell lines and on 505 clinical tumor specimens. JuLI has several characteristics. First, with the implementation of the noise reduction algorithm to minimize false positive calls, it maintains good analytical specificity without loss of sensitivity, particularly in noisy samples, such as FFPE samples. Second, JuLI can detect fusions with insufficient evidence in serial or multi-region sequencing samples by using the joint call function. Third, JuLI is easy to use with the provided an R package, which is available via GitHub (https://github.com/sgilab/JuLI) and supports comprehensive annotation and visualization of SVs.

An intriguing point is that JuLI can be used for monitoring cancer in specimens such as cfDNA, blood samples of minimal residual leukemic cell follow-up, and follow-up biopsy specimen without ideal tumor purity (Shin et al. 2017). Split reads originating from the primary tumor have high specificity in the location of fusion junction and adjacent DNA sequences, with uniqueness of split portion. Sensitive calling for these predefined fusion signals in follow-up specimens provided a good chance for early detection of relapsing cancer with good specificity.

Clinically, quantitative evaluation of fusion transcript is emphasized in follow-up of some cancers, particularly for CML. Discontinuation of TKIs is suggested in recent NCCN guidelines for CML, and the practice is performed based on quantitative evaluation of the *BCR-ABL1* transcript. Although the normalization, RNA assay does not provide direct information on the number of malignant clones. However, ultra-high coverage next-generation sequencing (NGS) for DNA fusion could provide direct information on the number of remnant malignant clones in future precision medicine.

For accurate detection of tumor-driving fusions, it is assumed to be necessary to detect fusions in both RNA and DNA. RNA is a suitable material for directly detecting chimeric transcripts; however, quality may be compromised because of long storage time or degradation during FFPE preparation (Ludyga et al. 2012). Detection sensitivity for fusions may be maximized by simultaneously performing DNA and RNA assays. Furthermore, combined fusion analysis for DNA and RNA can help identify loss-of-function of a tumor suppressor gene via fusion or complex fusions involving noncoding regions.

The limitation of this study is that a limited number of fusion events were tested for performance validation. Most callers used in this study for comparison reported their results through a genome-wide comparison of several samples in their papers. However, we performed quality (i.e., condition)-wide comparison of 505 patient samples with three clinically important fusion events and four cancer cell lines with known fusions. In a clinical setting, it may be inevitable to examine tissues with inappropriate quality (low tumor purity or poor quality FFPE). Therefore, our results could provide useful information to select callers in a clinical setting.

In clinical setting, although sensitivity is important, maintaining PPV is also essential to reduce the number of false positives. In particular, if the prevalence is relatively low, such as that of the *ALK* fusion in NSCLC (2%–7%) (Kwak et al. 2010), several wrong decisions can be made when PPV is not guaranteed. Clinical decisions based on false test results are risky and may lead to inappropriate treatment strategies. Therefore, if the PPV cannot provide a sufficiently high confidence level, it will be difficult to use the method for diagnostic purposes. Because JuLI has better PPV relative to the existing algorithms, it is likely to deliver accurate fusion profiling data to help clinicians to make optimal therapeutic decisions.

## CODE AVAILABILITY

The R package of JuLI is available online at GitHub (https://github.com/sgilab/JuLI).

## DATA AVAILABILITY

The sequence data of cell lines have been deposited in NCBI sequence read archive (SRA) under accession number (PRJNA514104). The patient data supporting the findings of this study are available on request from the corresponding author [W.-Y.P]. The raw data on patients are not publicly available because we did not have explicit consent to share the raw data acquired from the collected samples for the clinical test.

## ACKNOWLEDGEMENTS

This study was part of H.-T.S’s doctoral dissertation. We thank Jungwook Park and Geunhan Jeong for their critical advice on the tool development.

## AUTHOR CONTRIBUTIONS

H.-T.S., S.K., and K.-W.L. performed bioinformatic analysis of all data, with guidance from W.-Y.P. The manuscript was written by H.-T.S., N.K.D.K., and J.W.Y. with substantial input from J.K., J.S.B., and D.P‥ H.-T.S., N.K.D.K., B.L., and D.R. developed the tool. Y.-L.C., S.-H.L., M.-J.A., and K.P. provided clinical data. All authors reviewed the manuscript.

## REFERENCES

Apweiler R, Bairoch A, Wu CH, Barker WC, Boeckmann B, Ferro S, Gasteiger E, Huang H, Lopez R, Magrane M et al. 2004. UniProt: the Universal Protein knowledgebase. Nucleic acids research 32(Database issue): D115–119.

Araujo LH, Timmers C, Shilo K, Zhao W, Zhang J, Yu L, Natarajan TG, Miller CJ, Yilmaz AS, Liu T et al. 2015. Impact of Pre-Analytical Variables on Cancer Targeted Gene Sequencing Efficiency. PloS one 10(11): e0143092.

Awad MM, Shaw AT. 2014. ALK inhibitors in non-small cell lung cancer: crizotinib and beyond. Clinical advances in hematology & oncology: H&O 12(7): 429–439.

Barros-Silva JD, Paulo P, Bakken AC, Cerveira N, Lovf M, Henrique R, Jeronimo C, Lothe RA, Skotheim RI, Teixeira MR. 2013. Novel 5’ fusion partners of ETV1 and ETV4 in prostate cancer. Neoplasia 15(7): 720–726.

Cancer Genome Atlas Research N. 2015. The Molecular Taxonomy of Primary Prostate Cancer. Cell 163(4): 1011–1025.

Chen X, Schulz-Trieglaff O, Shaw R, Barnes B, Schlesinger F, Kallberg M, Cox AJ, Kruglyak S, Saunders CT. 2016. Manta: rapid detection of structural variants and indels for germline and cancer sequencing applications. Bioinformatics 32(8): 1220–1222.

Chong Z, Ruan J, Gao M, Zhou W, Chen T, Fan X, Ding L, Lee AY, Boutros P, Chen J et al. 2017. novoBreak: local assembly for breakpoint detection in cancer genomes. Nature methods 14(1): 65–67.

Christensen E, Nordentoft I, Vang S, Birkenkamp-Demtroder K, Jensen JB, Agerbaek M, Pedersen JS, Dyrskjot L. 2018. Optimized targeted sequencing of cell-free plasma DNA from bladder cancer patients. Scientific reports 8(1): 1917.

Dagogo-Jack I, Brannon AR, Ferris LA, Campbell CD, Lin JJ, Schultz KR, Ackil J, Stevens S, Dardaei L, Yoda S et al. 2018. Tracking the Evolution of Resistance to ALK Tyrosine Kinase Inhibitors through Longitudinal Analysis of Circulating Tumor DNA. JCO precision oncology 2018.

Druker BJ, Talpaz M, Resta DJ, Peng B, Buchdunger E, Ford JM, Lydon NB, Kantarjian H, Capdeville R, Ohno-Jones S et al. 2001. Efficacy and safety of a specific inhibitor of the BCR-ABL tyrosine kinase in chronic myeloid leukemia. The New England journal of medicine 344(14): 1031–1037.

Evers DL, He J, Kim YH, Mason JT, O’Leary TJ. 2011. Paraffin embedding contributes to RNA aggregation, reduced RNA yield, and low RNA quality. The Journal of molecular diagnostics: JMD 13(6): 687–694.

Finn RD, Mistry J, Tate J, Coggill P, Heger A, Pollington JE, Gavin OL, Gunasekaran P, Ceric G, Forslund K et al. 2010. The Pfam protein families database. Nucleic acids research 38(Database issue): D211–222.

Forbes SA, Beare D, Gunasekaran P, Leung K, Bindal N, Boutselakis H, Ding M, Bamford S, Cole C, Ward S et al. 2015. COSMIC: exploring the world’s knowledge of somatic mutations in human cancer. Nucleic acids research 43(Database issue): D805–811.

Frampton GM, Fichtenholtz A, Otto GA, Wang K, Downing SR, He J, Schnall-Levin M, White J, Sanford EM, An P et al. 2013. Development and validation of a clinical cancer genomic profiling test based on massively parallel DNA sequencing. Nature biotechnology 31(11): 1023–1031.

H. l. 2011. wgsim - Read simulator for next generation sequencing. Github Repository.

Hallberg B, Palmer RH. 2013. Mechanistic insight into ALK receptor tyrosine kinase in human cancer biology. Nature reviews Cancer 13(10): 685–700.

Kent WJ, Sugnet CW, Furey TS, Roskin KM, Pringle TH, Zahler AM, Haussler D. 2002. The human genome browser at UCSC. Genome research 12(6): 996–1006.

Kwak EL, Bang YJ, Camidge DR, Shaw AT, Solomon B, Maki RG, Ou SH, Dezube BJ, Janne PA, Costa DB et al. 2010. Anaplastic lymphoma kinase inhibition in non-small-cell lung cancer. The New England journal of medicine 363(18): 1693–1703.

Layer RM, Chiang C, Quinlan AR, Hall IM. 2014. LUMPY: a probabilistic framework for structural variant discovery. Genome biology 15(6): R84.

Lee SE, Lee B, Hong M, Song JY, Jung K, Lira ME, Mao M, Han J, Kim J, Choi YL. 2015. Comprehensive analysis of RET and ROS1 rearrangement in lung adenocarcinoma. Modern pathology: an official journal of the United States and Canadian Academy of Pathology, Inc 28(4): 468–479.

Li H, Durbin R. 2009. Fast and accurate short read alignment with Burrows-Wheeler transform. Bioinformatics 25(14): 1754–1760.

Li H, Handsaker B, Wysoker A, Fennell T, Ruan J, Homer N, Marth G, Abecasis G, Durbin R, Genome Project Data Processing S. 2009. The Sequence Alignment/Map format and SAMtools. Bioinformatics 25(16): 2078–2079.

Ludyga N, Grunwald B, Azimzadeh O, Englert S, Hofler H, Tapio S, Aubele M. 2012. Nucleic acids from long-term preserved FFPE tissues are suitable for downstream analyses. Virchows Archiv: an international journal of pathology 460(2): 131–140.

McKenna A, Hanna M, Banks E, Sivachenko A, Cibulskis K, Kernytsky A, Garimella K, Altshuler D, Gabriel S, Daly M et al. 2010. The Genome Analysis Toolkit: a MapReduce framework for analyzing next-generation DNA sequencing data. Genome research 20(9): 1297–1303.

Noh KW, Lee MS, Lee SE, Song JY, Shin HT, Kim YJ, Oh DY, Jung K, Sung M, Kim M et al. 2017. Molecular breakdown: a comprehensive view of anaplastic lymphoma kinase (ALK)-rearranged non-small cell lung cancer. The Journal of pathology 243(3): 307–319.

Oellerich M, Schutz E, Beck J, Kanzow P, Plowman PN, Weiss GJ, Walson PD. 2017. Using circulating cell-free DNA to monitor personalized cancer therapy. Critical reviews in clinical laboratory sciences 54(3): 205–218.

Pan Y, Zhang Y, Li Y, Hu H, Wang L, Li H, Wang R, Ye T, Luo X, Zhang Y et al. 2014. ALK, ROS1 and RET fusions in 1139 lung adenocarcinomas: a comprehensive study of common and fusion pattern-specific clinicopathologic, histologic and cytologic features. Lung cancer 84(2): 121–126.

Park G, Park JK, Son DS, Shin SH, Kim YJ, Jeon HJ, Lee J, Park WY, Lee KH, Park D. 2018. Utility of targeted deep sequencing for detecting circulating tumor DNA in pancreatic cancer patients. Scientific reports 8(1): 11631.

Paweletz CP, Sacher AG, Raymond CK, Alden RS, O’Connell A, Mach SL, Kuang Y, Gandhi L, Kirschmeier P, English JM et al. 2016. Bias-Corrected Targeted Next-Generation Sequencing for Rapid, Multiplexed Detection of Actionable Alterations in Cell-Free DNA from Advanced Lung Cancer Patients. Clinical cancer research: an official journal of the American Association for Cancer Research 22(4): 915–922.

Phallen J, Sausen M, Adleff V, Leal A, Hruban C, White J, Anagnostou V, Fiksel J, Cristiano S, Papp E et al. 2017. Direct detection of early-stage cancers using circulating tumor DNA. Science translational medicine 9(403).

Rausch T, Zichner T, Schlattl A, Stutz AM, Benes V, Korbel JO. 2012. DELLY: structural variant discovery by integrated paired-end and split-read analysis. Bioinformatics 28(18): i333–i339.

Shaw AT, Kim DW, Nakagawa K, Seto T, Crino L, Ahn MJ, De Pas T, Besse B, Solomon BJ, Blackhall F et al. 2013. Crizotinib versus chemotherapy in advanced ALK-positive lung cancer. The New England journal of medicine 368(25): 2385–2394.

Shin HT, Choi YL, Yun JW, Kim NKD, Kim SY, Jeon HJ, Nam JY, Lee C, Ryu D, Kim SC et al. 2017. Prevalence and detection of low-allele-fraction variants in clinical cancer samples. Nature communications 8(1): 1377.

Spencer DH, Sehn JK, Abel HJ, Watson MA, Pfeifer JD, Duncavage EJ. 2013. Comparison of clinical targeted next-generation sequence data from formalin-fixed and fresh-frozen tissue specimens. The Journal of molecular diagnostics: JMD 15(5): 623–633.

Takeuchi K, Soda M, Togashi Y, Suzuki R, Sakata S, Hatano S, Asaka R, Hamanaka W, Ninomiya H, Uehara H et al. 2012. RET, ROS1 and ALK fusions in lung cancer. Nature medicine 18(3): 378–381.

Thompson JC, Yee SS, Troxel AB, Savitch SL, Fan R, Balli D, Lieberman DB, Morrissette JD, Evans TL, Bauml J et al. 2016. Detection of Therapeutically Targetable Driver and Resistance Mutations in Lung Cancer Patients by Next-Generation Sequencing of Cell-Free Circulating Tumor DNA. Clinical cancer research: an official journal of the American Association for Cancer Research 22(23): 5772–5782.

Wala JA, Bandopadhayay P, Greenwald NF, O’Rourke R, Sharpe T, Stewart C, Schumacher S, Li Y, Weischenfeldt J, Yao X et al. 2018. SvABA: genome-wide detection of structural variants and indels by local assembly. Genome research 28(4): 581–591.

Zhou W, Chen T, Zhao H, Eterovic AK, Meric-Bernstam F, Mills GB, Chen K. 2014. Bias from removing read duplication in ultra-deep sequencing experiments. Bioinformatics 30(8): 1073–1080.

